# GenTB: A user-friendly genome-based predictor for tuberculosis resistance powered by machine learning

**DOI:** 10.1101/2021.03.27.437319

**Authors:** Matthias I Gröschel, Martin Owens, Luca Freschi, Roger Vargas, Maximilian G Marin, Jody Phelan, Zamin Iqbal, Avika Dixit, Maha R Farhat

## Abstract

**Introduction:** Multidrug-resistant *Mycobacterium tuberculosis* (*Mtb*) is a significant global public health threat. Genotypic resistance prediction from *Mtb* DNA sequences offers an alternative to laboratory-based drug-susceptibility testing. User-friendly and accurate resistance prediction tools are needed to enable public health and clinical practitioners to rapidly diagnose resistance and inform treatment regimens.

**Methods:** We present Translational Genomics platform for Tuberculosis (GenTB), a web-based application to predict antibiotic resistance from next-generation sequence data. The user can choose between two potential predictors, a Random Forest (RF) classifier and a Wide and Deep Neural Network (WDNN) to predict phenotypic resistance to 13 and 10 anti-tuberculosis drugs, respectively. We benchmark GenTB’s predictive performance along with leading TB resistance prediction tools (Mykrobe and TB-Profiler) using a ground truth dataset of 20,408 isolates with laboratory-based drug susceptibility data.

**Results:** All four tools reliably predicted resistance to first-line tuberculosis drugs but had varying performance for second-line drugs. The mean sensitivities for GenTB-RF and GenTB-WDNN across the nine shared drugs was 77.6% (95% CI 76.6 - 78.5%) and 75.4% (95% CI 74.5 - 76.4%) respectively, and marginally higher than the sensitivities of TB-Profiler at 74.4% (95% CI 73.4 - 75.3%) and Mykrobe at 71.9% (95% CI 70.9 - 72.9%). The higher sensitivities were at an expense of ≤1.5% lower specificity: Mykrobe 97.6% (95% CI 97.5 - 97.7%), TB-Profiler 96.9% (95% CI 96.7 to 97.0%), GenTB-WDNN 96.2% (95% CI 96.0 to 96.4%), and GenTB-RF 96.1% (95% CI 96.0 to 96.3%). Genotypic resistance sensitivity was 11% and 9% lower for isoniazid and rifampicin respectively, on isolates sequenced at low depth (<10x across 95% of the genome) emphasizing the need to quality control input sequence data before prediction. We discuss differences between tools in reporting results to the user including variants underlying the resistance calls and any novel or indeterminate variants

**Conclusion:** GenTB is an easy-to-use online tool to rapidly and accurately predict resistance to anti-tuberculosis drugs. GenTB can be accessed online at https://gentb.hms.harvard.edu, and the source code is available at https://github.com/farhat-lab/gentb-site.

## INTRODUCTION

Human tuberculosis, a chronic infectious disease caused by members of the *Mycobacterium tuberculosis* complex, is a leading cause of death from a bacterial infectious agent [1]. The proliferation of multidrug-resistant tuberculosis (MDR-TB) is threatening TB prevention and control activities worldwide [1]. Timely detection of antimicrobial resistance is vital to guide therapeutic options and contain transmission. Antimicrobial resistance is conventionally determined by *in vitro* drug susceptibility tests (DST) on solid or liquid antibiotic-containing culture, which uses drug-specific testing breakpoints (‘critical concentration’) to classify the infecting strain into drug-susceptible or drug-resistant [2]. Being contingent on mycobacteria’s slow growth rate, these phenotypic tests require days to weeks and often deliver unreliable and poorly reproducible results for some drugs, such as ethambutol and pyrazinamide [3,4]. In contrast, molecular methods have emerged as rapid resistance prediction alternatives to complement and speed up traditional DST, leveraging known and reliable genotype-phenotype relationships between variants in the *M. tuberculosis* genome and *in vitro* drug resistance [5].

Over recent years, whole-genome sequencing (WGS) of *M. tuberculosis* has become an affordable tool to provide genetic information for genotypic resistance prediction and high-resolution outbreak reconstruction [6]. Large scale genotype-phenotype assessments have demonstrated high diagnostic accuracy for clinical use to predict susceptibility to first-line drugs based on WGS [7]. Following these results, public health authorities have begun to discontinue phenotypic testing when pan susceptibility is predicted from the genotype, a step with considerable cost- and time benefits [8]. Start-to-end applications which analyze sequencing data to predict resistance phenotypes and are accessible to non-bioinformatic experts are required as WGS based analyses become part of the standardized diagnostic process in clinical laboratories. A range of published tools available for command-line [9,10] or web-based/desktop use [11–13] or both [14,15] exists. These applications vary in quality control and sequence preprocessing steps and rely on detecting pre-defined resistance-conferring mutations such as single nucleotide polymorphisms (SNPs) or small insertions/deletions (indels) in the WGS data to predict the resistance phenotype. They also vary in the type of information fed back to the user including error rates and specific variants detected.

Here, we present GenTB (https://gentb.hms.harvard.edu), an open user-friendly start-to-end application to predict drug resistance phenotypes to 13 drugs from WGS data. The GenTB analysis pipeline is also available for command-line use wrapped in *Snakemake* [16]. The online user interface allows users to interactively explore the sequencing data, prediction results and geographic distributions. Resistance prediction is made based on a previously observed set of variant positions spanning 18 resistance-associated genetic loci and a validated random forest (RF) classifier [17] as well as a wide and deep neural network (WDNN) combining a logistic regression model with a multilayer perceptron to predict the resistance phenotype [18]. In this study, we benchmark these two classification models implemented in GenTB along with two other tools with a command-line interface, *TB-profiler*, and *Mykrobe*, on a large dataset of >20k clinical *M. tuberculosis* isolates starting from raw Illumina sequence data.

## METHODS

### Backend and website build

GenTB is a bespoke Django website hosted by the Harvard Medical School O2 high performance computing environment and collaboratively developed on GitHub (https://github.com/farhat-lab/gentb-site). The website uses off-the-shelf frontend components; Bootstrap for styling and mobile-friendly delivery, nvd3 for plots and graphs, resumable.js for robust uploading and supplements these with custom Javascript functionality for integration. The backend is a Python-Django web service using a PostgreSQL database which integrates with Dropbox for file uploading, and python-chore for slurm cluster job submission and management. GenTB predict jobs are run by modular programs organized into pipelines. The modularity allows for easy maintenance and management of dependencies and outputs. Administration screens allow a non-expert developer design new program calls and construct new pipelines and integrate them without redeployment of the website. Further tools provide error tracking. GenTB predict results are integrated into the PostgreSQL database allowing website generated plots to be populated quickly. All generated files for the intermediary pipeline steps are provided for download by the user. GenTB Map uses a PostGIS database to rapidly link strain mutation and lineage information with geo-spatial objects; these are fed into the leaflet.js display to render strain information to the user. Map allows users to display strain data groupings by country, lineage, drug resistance phenotype or specific genetic mutation through tabs that can nest the groups in any order.

### Raw read processing

Upon uploading single-end or paired-end FastQ files, GenTB first validates the input using *fastQValidator* (Fig. 1). Low-quality reads and sequencing adapters are then trimmed with *fastp* [19]. Read mapping taxonomy is assessed with a custom-built *Kraken* database comprising *M. tuberculosis* complex reference sequences [20] followed by *minimap2* alignment (parameters: default) of reads to the H37Rv reference genome (AL123456) [21]. *Samtools* is used for sorting the aligned reads, removing duplicates, and indexing [22]. Sequence read datasets with a coverage of <95% at 10x or less across the genome or that had a mapping percentage of <90% to *M. tuberculosis* complex strains will not be further processed, and an error message is displayed to the user. Variants are called with *pilon* (parameters: default) [23] to obtain SNPs and indels in the variant calling format (VCF) requiring that they have a PASS or Amb filter tags with read allele frequency >0.40. *Fast-Lineage-Caller* then detects the *M. tuberculosis* lineage based on five lineage typing schemes as implemented by Freschi *et al*. [24]. Subsequently, invariant sites in the VCF file are removed, and a custom Perl script annotates each variant as frameshift, synonymous or non-synonymous, stop codon, indel along with the H37Rv locus tag for each respective gene. A custom python script generates a matrix file with all model features/variables in the columns used as input to the two prediction steps specified below. These scripts are available from Github (https://github.com/farhat-lab/gentb-site) and are open source (AGPLv3 license). All intermediate sequence files are accessible to the user for download and verification.

**Figure 1.**
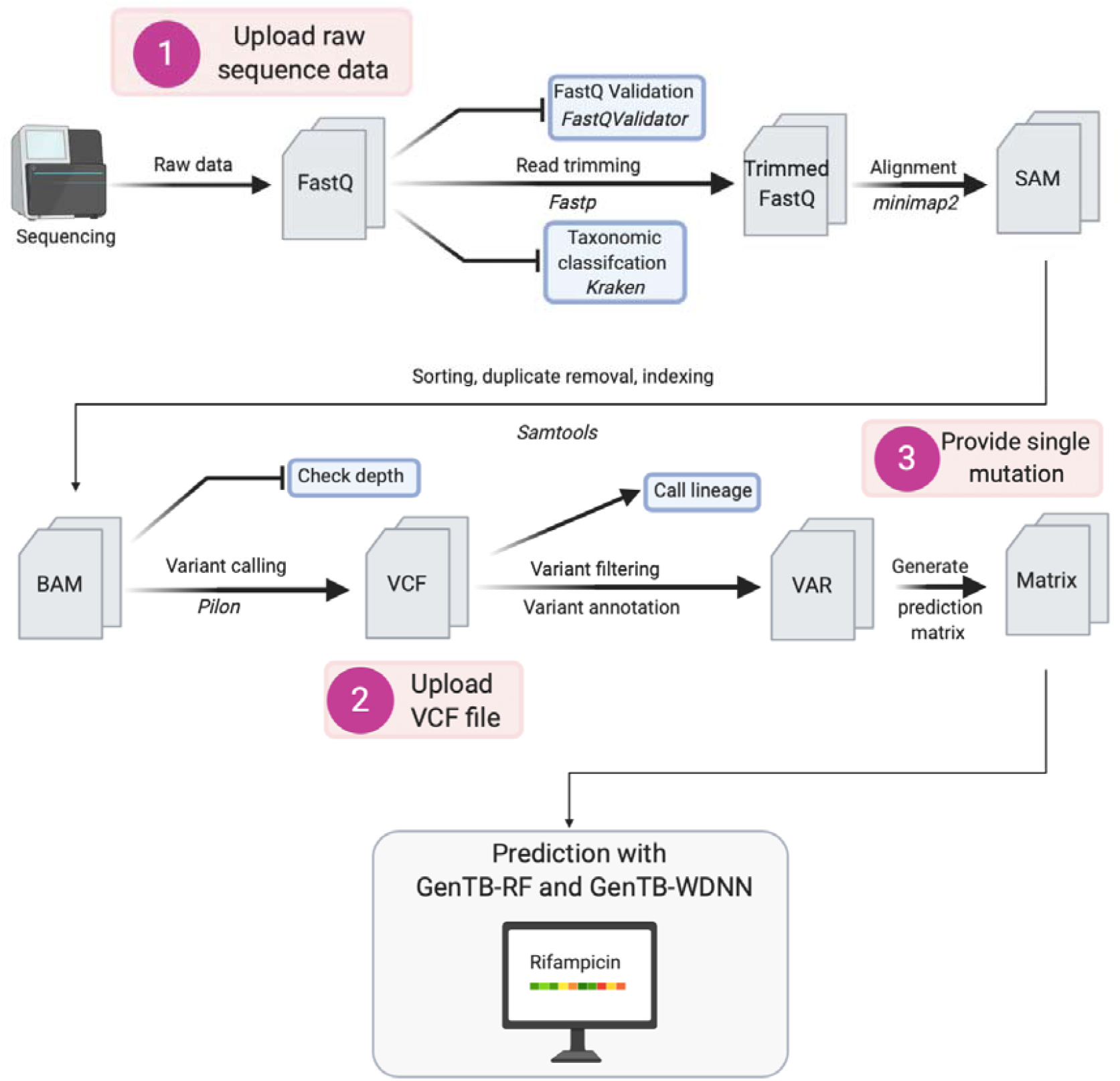
Schematic overview of the GenTB pipeline. Raw sequence data is quality checked and adapter trimmed before alignment to the H37Rv reference strain (accession AL123456). Variants are called with Pilon, and a variant matrix used by the prediction models are prepared using custom scripts available on Github. The analysis will fail if quality criteria are not met (blunt end arrows). Numbers represent the three moments in the pipeline where users can upload their data to predict resistance for their isolate.

### Operation

Users must create an account to run predictions and track uploaded datasets, intermediary files and results. Users with low internet bandwidth can use the *Dropbox* integration to upload files. Both raw sequence reads and variants in variant call format (VCF) can be uploaded for resistance prediction. The user can select an option to delete uploaded source data after prediction or otherwise to save it for their future access through GenTB. Files are user specific and not shared or accessible by others.

GenTB online interface has been tested with batches of up to 300 isolates. For batch processing of larger numbers of raw sequence data, we provide a command-line GenTB workflow based on *Snakemake* v5.20.1 [16] where dependent software will be sourced via conda [25]. The *Snakemake* workflow can be accessed via Github (https://github.com/farhat-lab/gentb-snakemake). The README file details how resistance prediction results on a paired-end sample can be obtained.

### Genotypic resistance prediction using two statistical models

Two multivariate models are used to predict the resistance phenotype, an RF model (GenTB-RF) and a WDNN (GenTB-WDNN). GenTB-RF was trained on isolates with available resistance phenotype data and was validated as previously described [17]. Briefly, 1,397 clinical isolates sampled as detailed in reference [17] underwent targeted sequencing at 18 drug resistance loci using molecular inversion probes and in parallel underwent binary drug culture-based DST to 13 drugs. One RF was built for each drug using the randomForest R package (v. 4.6.7) with a subset of the total 992 SNPs/indels observed. Variants of highest importance for resistance prediction to each drug were selected by iteratively paring down the model and measuring loss of performance. Important variants are shown in Suppl. Figure S1 for isoniazid and rifampicin.

Pyrazinamide resistance is known to rely on a large number of individually rare variants. Given the large increase in published *M. tuberculosis* WGS and linked DST data as well as the recent implication of novel resistance loci we retrained the pyrazinamide RF here using a newer version of randomForest R package (v. 4.6.-14) on variants in the genes *pncA, panD, clpC1, clpP* [26]. We used 75% (15,267 isolates) of the dataset to train the model and 25% (5,098 isolates) to validate its performance. During retraining, we excluded silent variants, those that occurred only in phenotypically susceptible isolates, or known phylogenetic variants, and the final model was trained on 393 variants occurring in 3,262 phenotypically pyrazinamide resistant isolates [24]. We chose the randomForest *mtry* variable that yielded the smallest out-of-bag error and varied the *classwt* variable to maximize the sum of sensitivity and specificity.

GenTB-WDNN is a multitask logistic regression model combined with a multilayer perceptron. It has been previously shown to have equal or higher performance than the RF architecture when both are trained on the same data [18]. GenTB-WDNN was trained on 3,601 isolates (sampled as detailed in reference [18]) for 11 drugs using the Keras 2.2.4 library in Python 3.6 with a TensorFlow 1.8.0 backend. The model uses 222 features (i.e., SNPs or small insertions/deletions) along with derived variables (i.e., the number of non-synonymous SNPs across all resistance-conferring genes) to predict the resistance phenotype.

### Validation sequencing and phenotype data

We collated a database of 20,408 Illumina raw sequence read datasets for which laboratory-based phenotypic DST data was available from public sources (Suppl. Table S1). Sequence data was downloaded from NCBI nucleotide databases. Custom scripts were used to pool the phenotype data from NCBI, Patric, ReseqTB, and the supplementary information from published literature (detailed methods in https://github.com/farhat-lab/resdata-ng). Sequence data was merged in case of multiple sequencing runs per isolate for downstream processing and resistance prediction. In isolates where >10% of reads did not classify as *M. tuberculosis* complex, we removed unclassified reads using seqtk (https://github.com/lh3/seqtk).

### Performance of GenTB and comparison with other tools

To assess the performance of GenTB for predicting resistance, all isolates were processed through the GenTB pipeline. We compared the diagnostic accuracy with two leading resistance prediction tools, *TB-profiler* 2.8.12 [14] and *Mykrobe* v0.9.0 [15], that were run with default parameters. The two tools and two GenTB prediction models’ predictive ability were obtained by comparing the genotypic prediction to the phenotype data that was considered the ground truth. We calculated the true positive rate (sensitivity), the true negative rate (specificity), and area under the receiver operating curve (AUC for short) to measure test accuracy for each drug and tool. We evaluated 1,000 probability thresholds per drug to call resistance or susceptibility for GenTB-RF while using the GenTB-WDNN thresholds previously described [18] (Suppl. Fig S2 and S3).

### Statistical Analyses and data visualization

Prediction files from all tools were parsed and analyzed in Jupyter Notebooks running Python 3.7 using the Pandas [27] and JSON libraries. Receiver operating characteristic curves were plotted using the Seaborn library [28]. The Vioplot package was used for violin plots [29]. Summary tables were created in R version 3.6.3 [30] using the packages from the tidyverse [31] and kable (https://cran.r-project.org/web/packages/kableExtra/index.html). Sequencing depth in resistance loci was calculated and plotted using *Mosdepth* version 0.2.9 [32]. Confidence intervals were obtained by bootstrapping, comparing 5000 predictions per tool and drug on a resampled dataset.

### Code and Data Availability

Code is available here: https://github.com/farhat-lab-gentb-site. The *snakemake* implementation is available here: https://github.com/farhat-lab/gentb-snakemake.

### Comparison of output between tools

We collated the output files and information produced by the GenTB online application, the webserver of TB-Profiler (https://tbdr.lshtm.ac.uk, version 3.0.0), and the Desktop version of Mykrobe (MacOS app v0.90) using one example raw sequence dataset (accession ERR1664619). The tools’ output was compared based on the following criteria: 1) Type and accessibility of output data formats; 2) Communication of genotypic prediction results, i.e. binary classification versus probability; 3) Disclosure of the prediction model’s error rate; 4) Description of known resistance conferring variants identified, 5) Reporting any novel mutation not listed in the resistance variant database, 6) Detailed account of detected lineage variants and what lineage typing scheme was used, 7) Report quality metrics on the input sequence data.

## RESULTS

### A user-friendly application to analyze *M. tuberculosis* sequencing data

GenTB was developed as a free and benchmarked online application to help public health and clinical practitioners deconvolute the complexity of *M. tuberculosis* WGS data. *GenTB Predict* allows users to predict resistance to 13 anti-TB drugs from a clinical isolate’s raw Illumina sequence data (FASTQ). Two validated machine learning models are used to make predictions: GenTB-RF and GenTB-WDNN (**Methods** and [17,18]). GenTB-RF is the default prediction model. In addition to the *GenTB Predict* function that we focus on here, the web-application has additional features for sharing, mapping, and exploring *M. tuberculosis* genetic and phenotypic data (Fig. 2). *GenTB Data* enables researchers to store, version, and share *M. tuberculosis* sequence and phenotype data and is powered by the Dataverse research data repository [33]. Users can select an option to delete source files upon processing the prediction. *GenTB Map* enables users to geographically visualize genetic and phenotype data. Users can explore the subset of 20,408 isolates with geographic tags (n= 12,547 isolates) used for GenTB predict validation (**Methods**), or can upload and explore their own data in enriched-VCF format (https://gitlab.com/doctormo/evcf/-/blob/master/docs/Enriched_VCF_Format.md). Raw data and results can be exported to a tabular data format.

**Figure 2.**
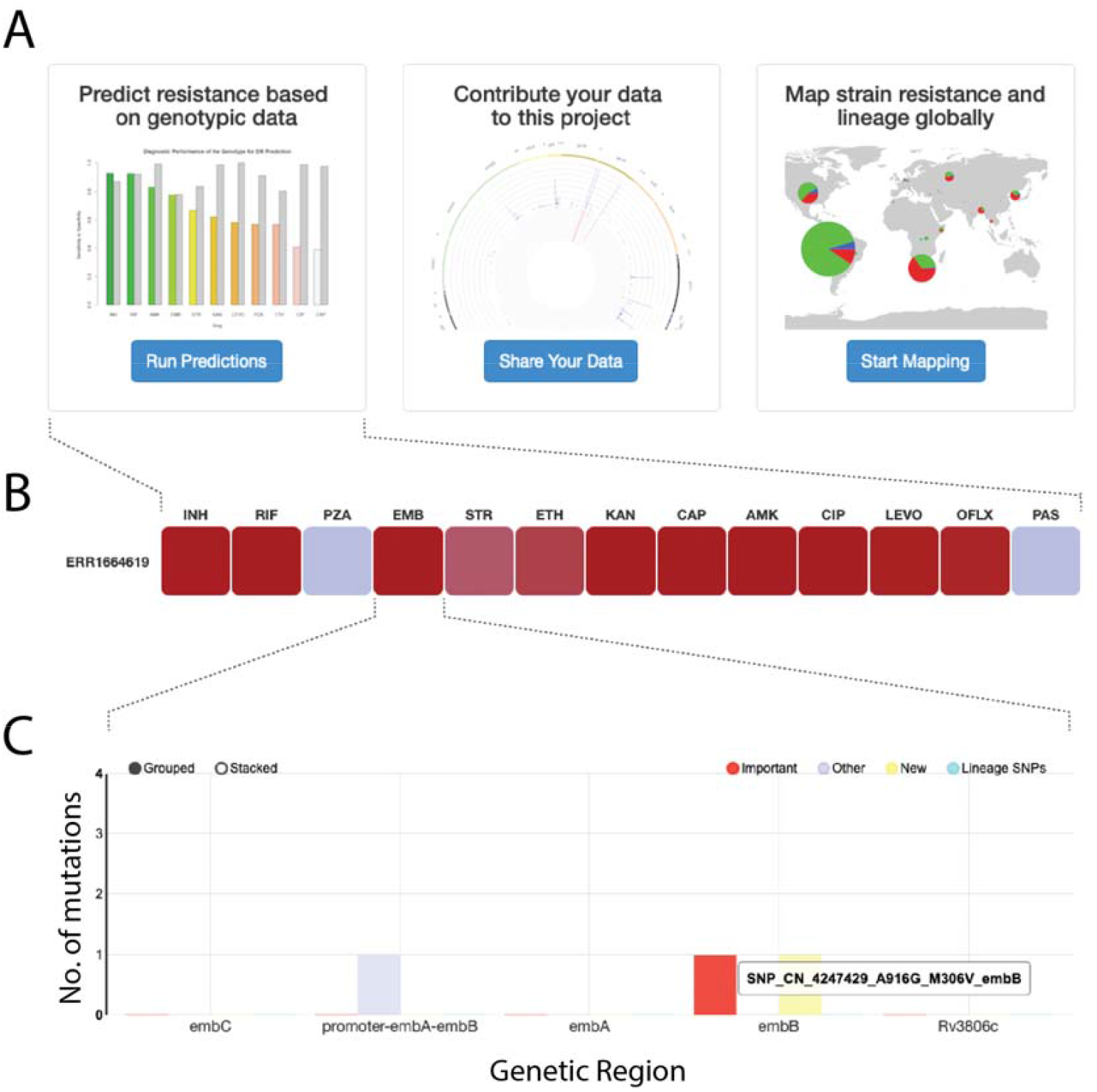
GenTB online user interface. **A)** The user is presented with the three main features offered by GenTB, i.e., to run predictions from user input data, to upload, share, and cite their data with the GenTB project, and to geographically map resistance frequencies or phenotype data. **B)** Example of a resistance prediction output where boxes are colored in the function of the prediction model’s output probability. **C)** Mutation plot that appears when clicked on one of the drugs heatmaps in **(B)**. Mutations will be shown when hovering the mouse over the genetic loci. INH = isoniazid, RIF = rifampicin, PZA = pyrazinamide, STR = streptomycin, EMB = ethambutol, ETH = ethionamide, KAN = kanamycin, CAP = capreomycin, AMK = amikacin, LEVO = levofloxacin, OFL = ofloxacin, PAS = Para-aminosalicylic acid.

### Dataset description

We curated a dataset of 20,408 *M. tuberculosis* isolates with known phenotypic resistance status to benchmark *GenTB Predict* performance (**Methods** and Suppl. Table S1). We excluded 29 isolates as they failed FastQ validation. Of the remaining, 1,339 isolates did not pass our taxonomy filter criterion, and their non-*M. tuberculosis complex* reads were removed. The GenTB pipeline identified an additional 499 isolates where more than 5% of the genome was covered at depth <10x and these isolates were excluded from further analysis. These isolates had a median depth of 21x (IQR 17 to 26). The remaining 19,880 isolates with high quality sequencing data were majority lineage 4 (52%), with lesser representation of lineage 2 (21%), lineage 3 (15%), lineage 1 (10%), *M. bovis* (0.6%), lineage 6 (0.3%), and lineage 5 (0.2%). Completeness of phenotypic DST data varied by drug and was highest for the first-line drugs rifampicin (98.3%), isoniazid (96.4%), ethambutol (77.5%), and pyrazinamide (71.5%) (Suppl. Table S2). The second and third-line drug phenotype data ranged from 35.1% completeness for streptomycin to 7.8% for ethionamide. Of the 20,408 isolates, 13,817 were phenotypically susceptible to first line drugs, 4,743 (23.3%) were phenotypically MDR (i.e., resistant to isoniazid and rifampicin) and 396 (1.9%) were phenotypically XDR (MDR and resistant to fluoroquinolones and the second-line injectables – amikacin, kanamycin or capreomycin). We ran GenTB-RF and GenTB-WDNN to predict resistance on 19,880 isolates and compared the predictions to phenotypic data.

### Predictive performance of the GenTB-Random Forest

We assessed each tools’ predictive performance by comparison with phenotypic culture-based DST results. Overall, the four tools had comparable performance characterized by varying sensitivities and high specificities (Tables 1 & 2, Fig 3A). Diagnostic performance was better for first-line than second-line drugs. As sensitivity varied most widely, we discuss it by drug class below. Specificities varied less by tool or by drug. GenTB-RF’s diagnostic specificity was >92% for all drugs including the second-line injectables and fluoroquinolones with the exception ethionamide (specificity = 78% [95% CI 75-80]) and streptomycin (specificity = 89% [95% CI 88-90]). GenTB-RF’s specificities were similar or higher than the other three tools with the exception of pyrazinamide (94% [95% CI 93-95]) and streptomycin (89% [95% CI = 88-90]) compared to TB-Profiler (96% and 95%, respectively) as well as Mykrobe (98% and 95%, respectively).

**Table 1:** Diagnostic accuracy of GenTB RandomForest and GenTB Wide and Deep Neural Network compared with two other leading prediction tools on a depth filtered dataset.

| DrugName | Phenotype |  | GenTB - RF |  | GenTB - WDNN |  | Mykrobe |  | TB-Profiler |  |
| --- | --- | --- | --- | --- | --- | --- | --- | --- | --- | --- |
| Isolates sequenced with high depth (n = 19.880) |  |  |  |  |  |  |  |  |  |  |
|  | R (n) | S (n) | Sensitivity (95% CI) | Specificity (95% CI) | Sensitivity (95% CI) | Specificity (95% CI) | Sensitivity (95% CI) | Specificity (95% CI) | Sensitivity (95% CI) | Specificity (95% CI) |
| isoniazid | 6,043 | 13,112 | 91% (91 to 92) | 98% (97 to 98) | 90% (89 to 91) | 99% (99 to 99) | 87% (86 to 88) | 98% (98 to 98) | 91% (90 to 92) | 98% (97 to 98) |
| rifampicin | 5,068 | 14,474 | 93% (93 to 94) | 98% (98 to 98) | 88% (88 to 89) | 99% (99 to 99) | 90% (89 to 91) | 98% (98 to 99) | 92% (91 to 93) | 98% (98 to 99) |
| ethambutol | 2,936 | 12,362 | 86% (85 to 87) | 92% (92 to 93) | 82% (80 to 83) | 93% (93 to 94) | 79% (77 to 80) | 93% (93 to 94) | 86% (85 to 88) | 92% (92 to 93) |
| pyrazinamide | 508 | 1,544 | 79% (76 to 83) | 94% (93 to 95) | 80% (79 to 82) | 95% (94 to 95) | 72% (71 to 74) | 98% (97 to 98) | 83% (80 to 86) | 96% (96 to 97) |
| amikacin | 618 | 3,458 | 67% (63 to 71) | 99% (99 to 100) | 66% (62 to 70) | 99% (99 to 100) | 63% (60 to 67) | 99% (99 to 100) | 55% (51 - 59) | 99% (99 to 100) |
| capreomycin | 648 | 3,733 | 63% (59 to 67) | 97% (97 to 98) | 57% (53 to 61) | 98% (98 to 99) | 60% (56 to 64) | 98% (98 to 99) | 56% (52 to 60) | 96% (95 to 96) |
| ethionamide | 502 | 1,094 | 67% (63 to 72) | 78% (75 to 80) | - | - | - | - | 70% (66 to 74) | 73% (70 to 76) |
| kanamycin | 576 | 3,707 | 68% (64 to 72) | 99% (98 to 99) | 66% (62 to 70) | 100% (99 to 100) | 66% (63 to 70) | 99 (99 to 100) | 68% (64 to 71) | 98% (98 to 99) |
| streptomycin | 2,126 | 4,968 | 82% (80 to 83) | 89% (88 to 90) | 87% (85 to 88) | 87% (86 to 88) | 68% (66 to 70) | 95% (95 to 96) | 71% (70 to 73) | 95% (95 to 96%) |
| ofloxacin | 743 | 4,038 | 68% (65 to 72) | 99% (98 to 99) | 62% (58 to 66) | 96% (95 to 96) | 62% (58 to 65) | 99% (98 to 99) | 67% (63 to 70) | 98 (98 to 99) |

**Figure 3:**
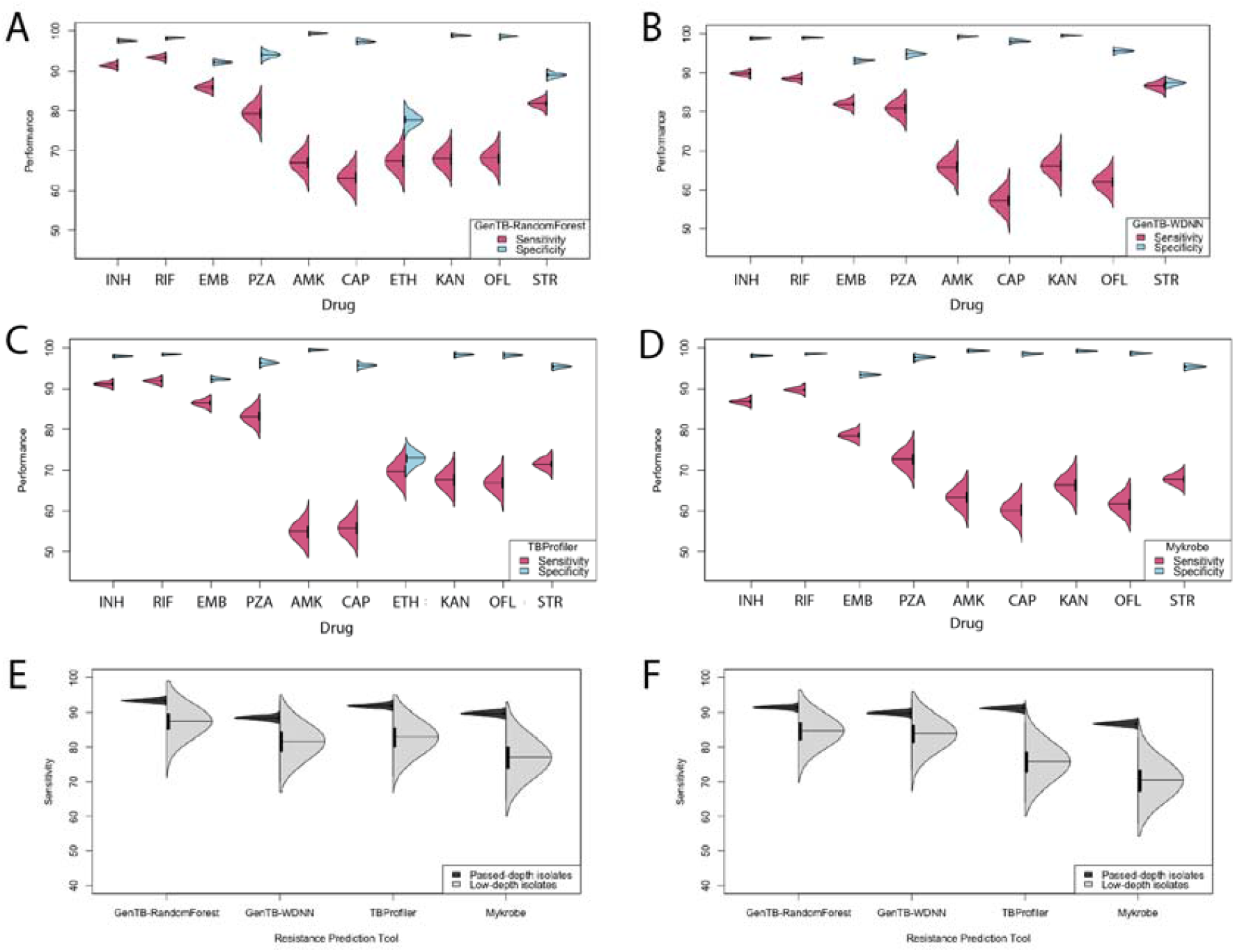
Diagnostic performance of the four prediction tools across antituberculosis drugs. Paired violin plots displaying sensitivity and specificity to predict drug resistance for **A)** GenTB-Random Forest, **B)** GenTB-Wide and Deep Neural Network, **C)** TB-Profiler and **D)** Mykrobe. **E)** Violinplot of diagnostic performance to predict rifampicin resistance comparing isolates passing depth filters (in black) to isolates that failed the depth-filters (in grey) arranged by prediction tool. **F)** Violinplot of diagnostic performance to predict isoniazid resistance comparing isolates passing depth filters (in black) to isolates that failed the depth-filters (in grey) arranged by prediction tool. AMK = amikacin, CAP = capreomycin, EMB = ethambutol, ETH = ethionamide, INH = isoniazid, KAN = kanamycin, OFL = ofloxacin, PZA = pyrazinamide, RIF = rifampicin, STR = streptomycin.

#### First-line drugs

Rifampicin resistance prediction by GenTB-RF was most accurate compared to other tools: AUC 0.96 (95% CI = 0.95-0.96), sensitivity 93% (95% CI = 93-94), second highest sensitivity was for TB-Profiler at 92% (95% CI = 91-93) (Tables 1 & 2, Figure 4). The accuracy of isoniazid resistance prediction was high and comparable across three of the four tools including GenTB-RF (sensitivity 91% [95% CI =91-92]). For ethambutol, GenTB-RF and TB-Profiler had the best and comparable performance with sensitivity 86% (95% CI =85-87).

**Figure 4:**
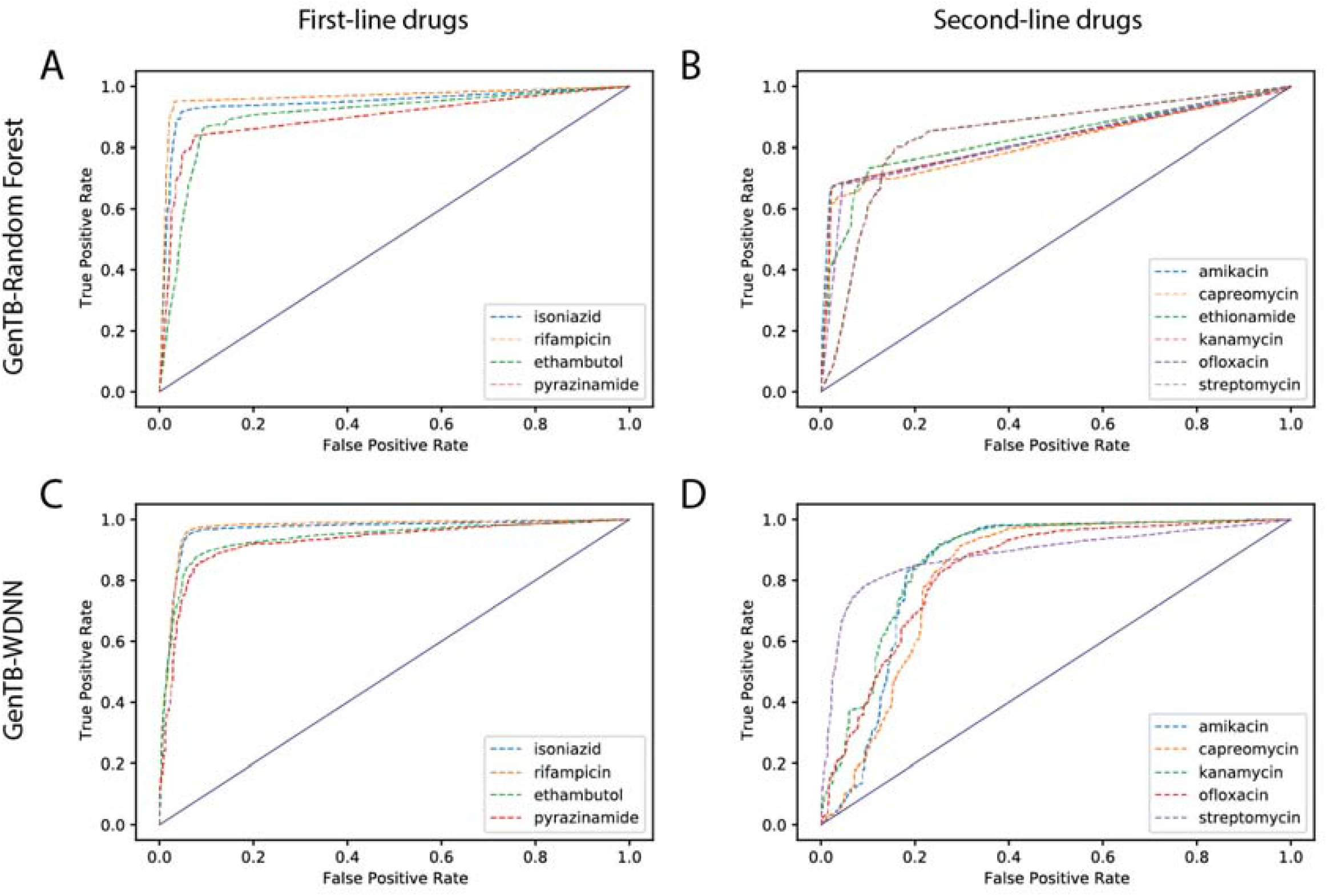
ROC performance curve of the GenTB-RF and GenTB-WDNN prediction models. A ROC plot of the GenTB-Random Forest (top) and GenTB-WDNN (bottom) predictive performance on the study dataset for first line (**A)** and **C)**) and second line drugs (**B)** and **D)**).

GenTB-RF predictions for pyrazinamide using the original model (v1.0) had low sensitivity at 56% (95% CI 54-58) with adequate specificity (98% [95% CI = 98-99]) compared to the other tools when evaluated on the 19,880 isolates (2,336 phenotypically resistant and 11,932 susceptible) [17]. Pyrazinamide resistance is known to be caused by a large number of individually rare variants in the gene *pncA* [34]. Given the large interval increase in available WGS data and recent implication of novel resistance loci (*panD, clpC1, clpP*) [26] since GenTB-RF was last trained, we assessed the number of rare variants in the four aforementioned genes linked to pyrazinamide resistance. In a random 75% subset of the 20,379 isolates, we detected a total of 393 different variants in *pncA, panD, clpC1 and clpP* with 40% (158/393) occurring only once. The majority of these variants, i.e., 73% (285/393) were not previously seen by the original model. As a result of these observations, we retrained a GenTB-RFv2.0, on 75% of the data using all 393 non-synonymous variants including singletons and insertion/deletion variants from *pncA, panD, clpC1* and *clpP*. The retrained model, when benchmarked on an independent validation dataset of 5,098 isolates, offered a sensitivity similar to that of the other tools (79%, 76 to 83) (Table 1).

#### Second-line drugs

For second-line drugs, larger discrepancies between genotype and resistance phenotype have been previously described compared with first-line drugs [14,15]. Resistance to the second-line injectable drugs amikacin and kanamycin ranged between 63-68% across the four tools, with the exception of a sensitivity of 55% by TB-Profiler for amikacin (Table 1). For the fluoroquinolone ofloxacin, sensitivity ranged from 62%-68% across the four tools. Three drugs had too few isolates with known phenotypic resistance (ciprofloxacin [n = 63], levofloxacin [n =111], and para-aminosalicylic acid [n = 46]), and hence the tool’s predictions had wide confidence intervals for these drugs (Supplemental Tables S3 and S4).

### Predictive performance of GenTB-WDNN

Similar to GenTB-RF, the overall GenTB-WDNN performance was marked by high prediction accuracy of first-line drug resistance and lower accuracy of second-line resistance (Table 1). AUC 95% CI overlapped for all drugs between the two models except for ofloxacin and rifampicin for which the GenTB-RF AUC was higher (Table 2). For streptomycin the GenTB-WDNN offered the best sensitivity and specificity of all four models (sensitivity 87%, 95% CI 85-88%, specificity 87% (95%CI 86-88%). Specificities were >95% for all drugs except for streptomycin (87%, 95% CI 85 to 88) and ethambutol (93%, 95% CI 93 to 94).

**Table 2:** Area under the Receiver Operating Characteristic curve for GenTB-RF and GenTB-WDNN

| Drug | GenTB-RF | GenTB-WDNN |
| --- | --- | --- |
|  | Area under the ROC curve (95% CI) |  |
| isoniazid | 0.94 (0.94 to 0.95) | 0.94 (0.94 to 0.95) |
| rifampicin | 0.96 (0.95 to 0.96) | 0.94 (0.93 to 0.94) |
| ethambutol | 0.89 (0.88 to 0.9) | 0.87 (0.87 to 0.87) |
| pyrazinamide | 0.90 (0.88 to 91) | 0.88 (0.87 to 0.88) |
| amikacin | 0.83 (0.81 to 0.85) | 0.83 (0.81 to 0.84) |
| capreomycin | 0.80 (0.78 to 0.82) | 0.78 (0.76 to 0.80) |
| ethionamide | 0.73 (0.7 to 0.75) | - |
| kanamycin | 0.83 (0.81 to 0.85) | 0.83 (0.81 to 0.85) |
| streptomycin | 0.85 (0.84 to 0.86) | 0.87 (0.86 to 0.88) |
| ofloxacin | 0.83 (0.82 to 0.85) | 0.79 (0.77 to 0.81) |

RF = Random Forest, WDNN = Wide and Deep Neural Network

### Predictive performance depends on sequencing depth

We evaluated the need for quality control on sequencing depth as several tools do not currently implement this prior to resistance prediction [9,14,15]. We observed predictive performance to be highly dependent on sequencing depth as indicated by lower sensitivity to predict rifampicin or isoniazid resistance by all four tools for the 499 isolates that did not meet the threshold of ≥10x depth across >95% of the genome (median depth of 21x, IQR 17 to 26, Figures 3E,3F). Using GenTB-RF, the mean sensitivity of isoniazid and rifampicin prediction was 84.6% (SD 3.6) and 87.3% (SD 3.6) respectively among low-depth isolates, compared with 91% and 93%, respectively, on high-depth isolates (Suppl. Table S5, Figures 3E, 3F). Loss of sensitivity due to low sequencing depth was comparable across the four tools.

### Discordant resistance predictions

To gain insight into model performance, we probed discrepancies between GenTB-RF’s genotype-based prediction and the resistance phenotype. We focused on this model as it had the highest overall sensitivity. We examined specifically rifampicin and isoniazid as resistance to these two drugs defines MDR-TB, and their genetic resistance mechanisms are well understood. We investigated isolates for which GenTB-RF predicted resistance while the phenotype was reported as susceptible (false positives) and isolates for which GenTB-RF predicted susceptibility with a resistant phenotype (false negatives). We confirmed that false negative predictions were not due to low sequencing depth in relevant drug resistance loci (*i*.*e*. that depth was ≥10x across all bases, Suppl. Figures S4 and S5).

#### Rifampicin false positives

Variants causative of rifampicin resistance are concentrated in a 81bp window in the *rpoB* gene *a*.*k*.*a* the rifampicin resistance determining region (RRDR, H37Rv coordinates 761081 to 761162, accession AL123456) [35]. For rifampicin, we observed 254 false positive predictions (phenotypically susceptible isolates predicted resistant). GenTB-RF detected one or more non-silent RRDR variants in 198 of these 254 isolates (78%). The most common RRDR variants were S450L (occurred in 49/254 isolates), L430P (in 33/254), and H445N (in 31/254) (Suppl. Table S6). The remaining 56 of 254 isolates, harbored non-RRDR variants, the two most common were *rpoB* I491F (occurred in 29/56) and *rpoB* V695L (occurred in 24/56). Twenty eight of the 56 isolates (50%) were phenotypically resistant to isoniazid and a further 16 (29%) were resistant to ethambutol.

#### Rifampicin false negatives

Among the 333 false negative rifampicin predictions (phenotypically resistant isolates predicted susceptible), 96 (29%) isolates harbored a variant in *rpoB* and of these 75 (23% of the 333) were in the RRDR (Suppl. Table S6). These included most commonly three base pair insertion in *rpoB* codon 433 (occurred in 14/333 isolates) and *rpoB codon* 443 (occurred in 9/333 isolates) and *rpoB* substitution Q432L (in 9/333) [36]. These *rpoB* variants were not previously seen by the GenTB-RF model when initially trained. For the remaining 237 of 333 isolates (71%) phenotypic resistance remained unexplained.

#### Isoniazid false positives

For isoniazid, we observed 315 false positive predictions (phenotypically susceptible isolates predicted resistant by GenTB-RF). Among these isolates, 119/315 (38%) had a total of 40 unique non-silent non-lineage variants in genes linked to isoniazid resistance (*inhA, katG, ahpC, fabG1*) (Suppl. Table S7). Most variants, 36/40, were rare, occurring in only 2 or fewer isolates. Five out of the 40 unique mutations detected in 75/315 (24%) isolates are considered important for isoniazid resistance prediction by GenTB-RF [17]. The most frequent INH resistance variants were the canonical isoniazid resistance mutation *katG* S315T [37] (occurred in 56/315 isolates) and non-silent variants at *inhA* codon 94 (occurred in 14/315 isolates). Seventy-six of the 315 (24%) apparent false positive isolates were phenotypically resistant to rifampicin and 189 (60%) isolates had a phenotypic resistance to at least one other drug.

#### Isoniazid false negatives

Among the 518 false negative isoniazid predictions (phenotypically resistant isolates predicted susceptible by GenTB-RF), 194/518 (37%) harbored non-silent variants in isoniazid resistance associated genes (Suppl. Table S7). Only 13 of the 139 unique variants observed in the 518 isolates were seen before by GenTB-RF and none of these were considered important isoniazid resistance mutations. *KatG* W328L was the variant detected most frequently (occurred in 10/518 isolates predicted false negative) and although not previously seen by GenTB-RF was described to occur in 0.2% of isoniazid resistance in one study [38]. Most variants linked to isoniazid resistance observed in these isolates were rare, i.e., 134/139 (96%) occurred in ≤ 3 isolates.

### Output comparison across the three tools

All four tools are accessible to the non-experienced user via either an online interface (GenTB, TB-Profiler) or via a Desktop application. We compared each tool’s output using the criteria specified in **Methods** (Table 3). GenTB-RF provides a heatmap indicating the probability of resistance including the models’ error rate with all prediction and intermediary files available for download. TB-Profiler and Mykrobe present binary (resistant or susceptible) predictions in overview tables with download options in CSV or JSON formats, respectively. TB-Profiler and GenTB present resistance causing variants and variants not associated with resistance. All tools provide the lineage call made but GenTB also specifies the lineage typing schemes used.

**Table 3:** Output comparison across tools

| Criteria | GenTB | TB-Profiler | Mykrobe |
| --- | --- | --- | --- |
| 1) Output |  |  |  |
| Type | Heatmap and barplot | Overview tables | Overview table |
| Download | all intermediate and output files (JSON) | yes (CSV) | Yes (JSON) |
| 2) Genotypic predictions | Probability | Binary | Binary |
| 3) Error rate | Yes | N.A. | N.A. |
| 4) Resistance variants | Variant by drug | Variant by drug<br>incl. fraction of mutant / wild-type allele | Variant by drug incl. depth of mutant and wild-type alleles |
| 5) Unknown variants | Yes, in all genes | Yes, in candidate resistance genes | No |
| 6) *M. tuberculosis* Lineage |  |  |  |
| Lineage | Yes | Yes | Yes |
| Typing scheme | Yes | No | No |
| 7) Quality metrics | Trimming and Kraken report downloadable | No. of reads,<br>Percentage of reads mapped | No |

## DISCUSSION

The increasing affordability of WGS and our improving comprehension of mycobacterial drug resistance mechanisms has placed sequencing at the forefront of *M. tuberculosis* resistance diagnosis in clinical and public health laboratories (*e*.*g*. Public Health England in the United Kingdom and the Centers for Disease Control and Prevention in the United States) [7,39]. Yet, the complexity of resistance biology is such that large and diverse bacterial isolate datasets are needed to confirm the accuracy of genotype-based resistance prediction and its generalizability. Further, the required computational resources and knowledge to conduct sequencing analysis prohibit both the access to and confidence in WGS based resistance prediction in clinics in both low- and high-incidence settings. High confidence automated tools that are systematically benchmarked on diverse datasets are needed to facilitate adoption, and to act as the standard for future tool development and regulation by oversight agencies such as the World Health Organization (WHO).

GenTB is an automated open tool for resistance prediction from WGS. Here we benchmarked its two prediction models against two other leading TB prediction tools. Both GenTB models predicted resistance and susceptibility against first-line drugs with high accuracy. Predictive performance for second line drugs showed lower sensitivity, although with high specificity for some of those drugs, i.e., capreomycin, kanamycin, and ofloxacin. This high specificity may be used to rule out resistance when no resistance conferring variant for these drugs was found. A detailed analysis of discrepant predictions made by GenTB-RF illustrated that a number of false positive predictions were supported by canonical resistance variants, e.g., non-silent mutation in the *rpoB* RRDR in case of rifampicin, suggesting that their phenotypes were erroneously labeled as susceptible. Similarly, nearly half (48%) of the variants found in isoniazid false positive predictions are canonical resistance variants. These isoniazid resistance variants, the large proportion (60%) of phenotypic resistance to another drug among these isolates, and the knowledge that isoniazid is usually a gateway drug resistance, suggest that some phenotypes were erroneously characterized as susceptible [40]. Accordingly, specificity of genotype-based prediction in practice maybe even higher than reported here (Table 1).

For isolates with a resistant rifampicin phenotype that were predicted susceptible by GenTB-RF, we found a mutation in the *rpoB* RRDR in a nearly a quarter (23%) of isolates that reasonably accounts for the resistance phenotype, but had not been seen by the model previously. For the remaining majority of false negatives (71% for rifampicin) no relevant resistance variant was found. In these cases, phenotypic resistance remained unexplained and could be due to erroneous phenotypes or yet unknown resistance mechanisms. For isolates with a resistant isoniazid phenotype predicted susceptible, no important resistance conferring mutations were found. In these cases, phenotypic resistance could be due to rare and yet undescribed resistance variants. A substantial proportion of false negative predictions to isoniazid or rifampicin had genotypic resistance to at least another drug (48% of rifampicin false negatives and 40% of isoniazid false negatives). These observations overall suggest that a viable option to reduce false negative predictions by current models would be to leverage genotypic predictions to other drugs and flag such isolates for complementary phenotypic DST. In the future as new larger datasets of paired genotype and resistance phenotype are curated, e.g. by efforts sponsored by the WHO [41], retraining existing resistance prediction models will improve diagnostic sensitivity.

The final output produced by the four tools varies in terms of detail and type of variants reported with GenTB providing the most detail. GenTB’s output reports novel variants not linked to resistance in addition to those that are resistance associated. The phylogenetic lineage calling procedure implemented in GenTB [24] uses currently available typing schemes, including the spoligotype nomenclature, to facilitate comparisons across lineage schemes.

Unlike other published resistance prediction tools that rely on a curated list of resistance conferring mutations that call resistance when a specific variant is present, GenTB-RF and GenTB-WDNN use multivariable statistical models to predict resistance phenotype. These models are better suited to account for the complex relationships between resistance genotype and phenotype. Among the advantages of multivariate prediction models is that relationships between variables are taken into account as both individual variants and gene-gene interactions cause phenotypic drug resistance. As such, the two models provide a probability value that a given isolate is resistant or susceptible rather than a binary classification. This is relevant in case of variants that, if present alone, confer only weak to no resistance, but may confer complete resistance if present in combination. Also, each variable in a multivariable model has different weights depending on the strength of association with resistance in the training data, reflecting the biological reality where variants cause differing levels of resistance. The benchmarking data presented here confirm that these multivariate models offer gains in sensitivity over the other two tools that use curated mutation lists, however this comes at a small decrease in specificity overall. Seen its higher overall performance GenTB-RF is currently implemented as the default prediction model. As larger and more diverse data will become available for model training, especially for prediction of resistance more quantitatively, i.e., to predict minimum inhibitory concentrations or MICs, we anticipate multivariate models including the more complex GenTB-WDNN architecture to have an even bigger advantage over direct association of mutation lists.

This study was not without limitations. An important prerequisite for reliable genotypic resistance prediction is the quality of the raw sequencing data. Variants and small indels in resistance conferring genes can be accurately and confidently called from Illumina raw sequence data if the genes are adequately covered at an acceptable sequencing depth [Marin *et al*., in preparation]. However, short-read sequencing data is recognized to have lower sensitivity for detecting more complex genomic variants including long indels or structural variation and these may have been missed in this study. But these latter types of variants are expected to be rare. Our finding of ‘apparent’ false positive predictions (i.e., resistance call by GenTB-RF while susceptible phenotype) in isolates harboring canonical resistance variants portends some erroneous phenotypes in our ground truth dataset. Due to the scale and public nature of the dataset used for benchmarking in this study, we were unable to retest the laboratory-based drug susceptibility profiles of isolates with discordant predictions, but hope that it provides a test closer to a ‘real-world’ scenario for these tool’s application.

## CONCLUSION

The rapid emergence and affordability of sequencing of *M. tuberculosis* along with the herein confirmed high accuracy of several genotypic resistance prediction tools supports the use of informatically assisted treatment design in the clinical setting. Independent benchmarking efforts will facilitate regulatory reviews and assessments and build confidence in the tools’ performances. As genotypic resistance predictions will accompany and increasingly replace laboratory-based resistance phenotyping performance criteria will need to be defined to guide clinical and public health laboratories in their use. Lastly, it will be important to communicate the confidence and uncertainty that is inherent to all genotypic predictions to clinicians, and provide clear diagnostic algorithms in case of genotype-phenotype discordances.

## Notes

### Competing Interest Statement

The authors have declared no competing interest.

### Summary of Updates

updated link to the GenTB websiet

