## Supplementary material for "GenTB: A user-friendly genome-based predictor for tuberculosis resistance powered by machine learning": Suppl. tables and figures

### SUPPLEMENTARY INFORMATION


**Authors:**

Matthias I Gröschel[1], Martin Owens[1], Luca Freschi[1], Roger Vargas Jr[1,2], Maximilian G Marin[1,2], Jody Phelan[3], Zamin Iqbal[4], Avika Dixit[1,5] and Maha R Farhat[1,6]

**Affiliations**

[1] Department of Biomedical Informatics, Harvard Medical School, Boston, MA, USA

[2] Department of Systems Biology, Harvard Medical School, Boston, MA, USA

[3] Faculty of Infectious and Tropical Diseases, London School of Hygiene & Tropical Medicine, London WC1E 7HT, UK

[4] European Bioinformatics Institute, Hinxton, Cambridge CB10 ISD, UK

[5] Division of Infectious Diseases, Boston Children's Hospital, Boston, MA, USA

[6] Division of Pulmonary and Critical Care Medicine, Massachusetts General Hospital, Boston, MA, USA

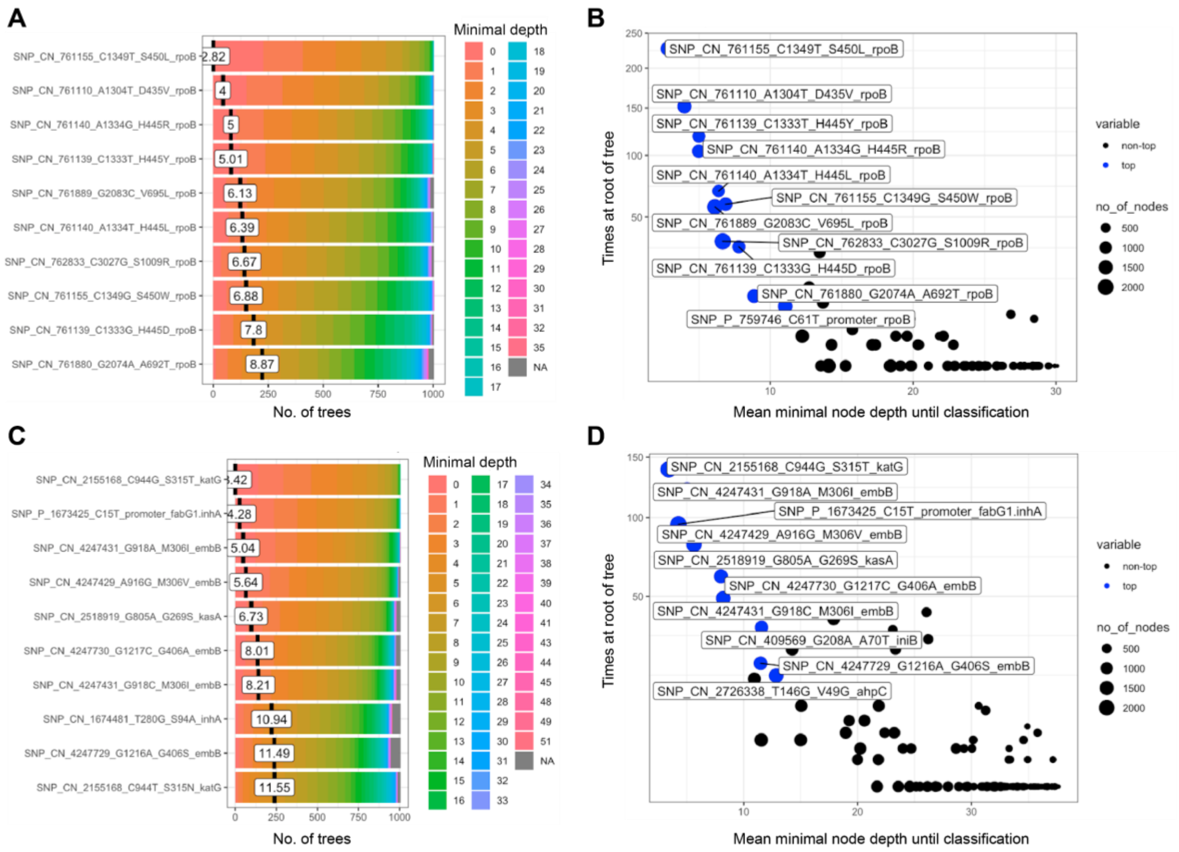


**Supplemental figure S1: Characteristics of variables used for resistance prediction by Gentb-RF for isoniazid and rifampicin. A)** Distribution of minimal node depth among the trees of the classification forest for rifampicin resistance is shown. The mean of the distribution is marked by a vertical bar with a value label on it, denoting the mean number of node depth required for the variant to reach classification into resistant or susceptible. **B)** Multi-way importance for rifampicin resistance classification showing mean depth of first split on the variant on the X-axis, the number of trees where the variable is at the root of the tree on the y-axis, and the total number of nodes in the forest that split on that variant by size of the dots. **C)** and **D)** as **A)** and **B)** for isoniazid resistance classification. Genetic variants are described as follows and are separated by underscores: First type of variant, second if the variant leads to change in amino acid (AA), frameshift, or stop codon, third the genomic coordinate based on the reference strain H37RV (AL123456), fourth AA change, fifth codon change, last locus tag.

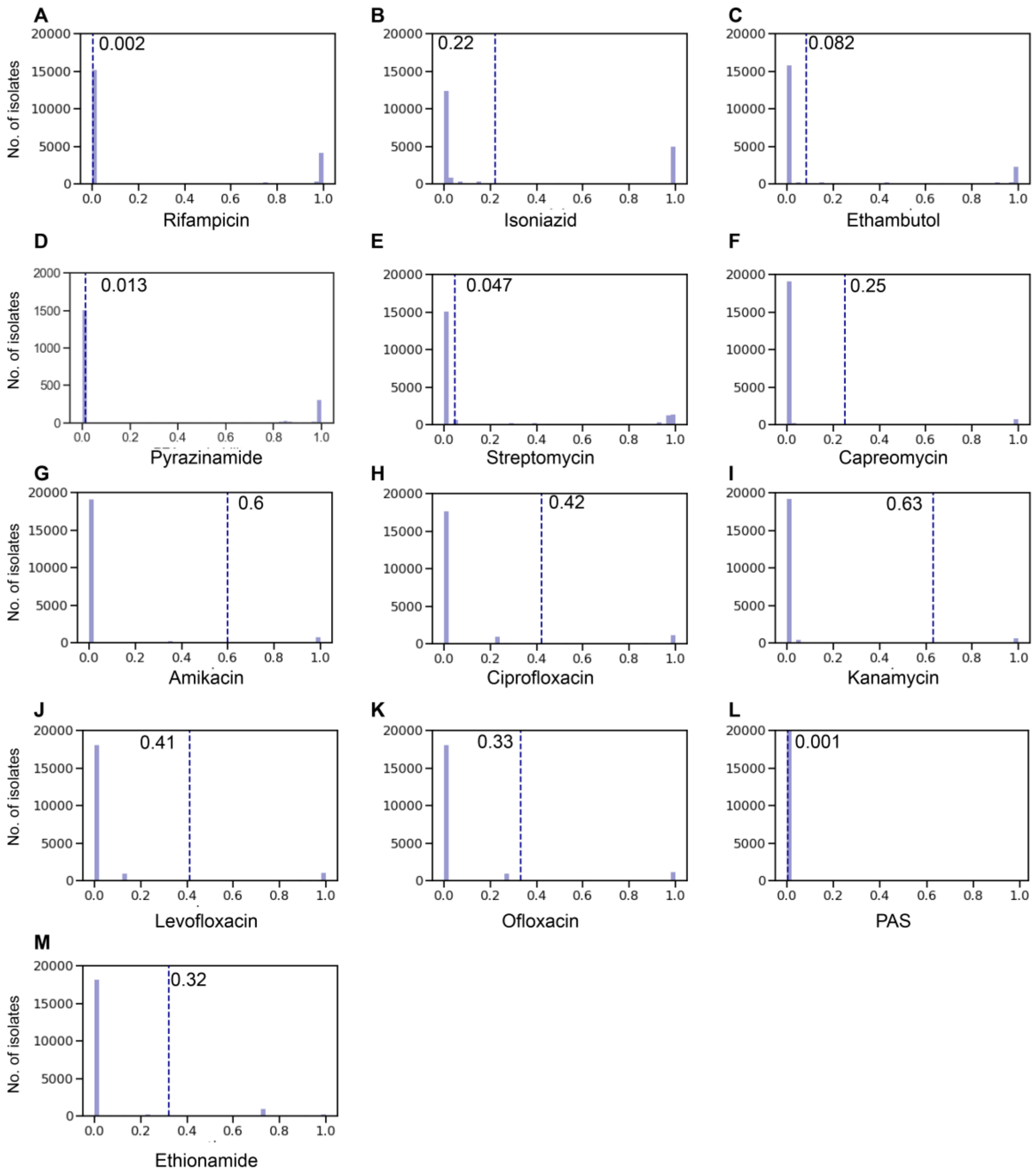


**Supplementary Figure S2: Probability distribution and selected threshold for Gentb-RF resistance predictions.** A-M. Probability of susceptibility (0 = drug susceptibility, 1 = drug resistance) for 13 drugs among the 20.379 isolates as predicted by Gentb-RF. Vertical lines depict the resistance threshold that yielded the highest predictive performance as measured by the sum of sensitivity and specificity. Drug-specific probability values are written in each subpanel.

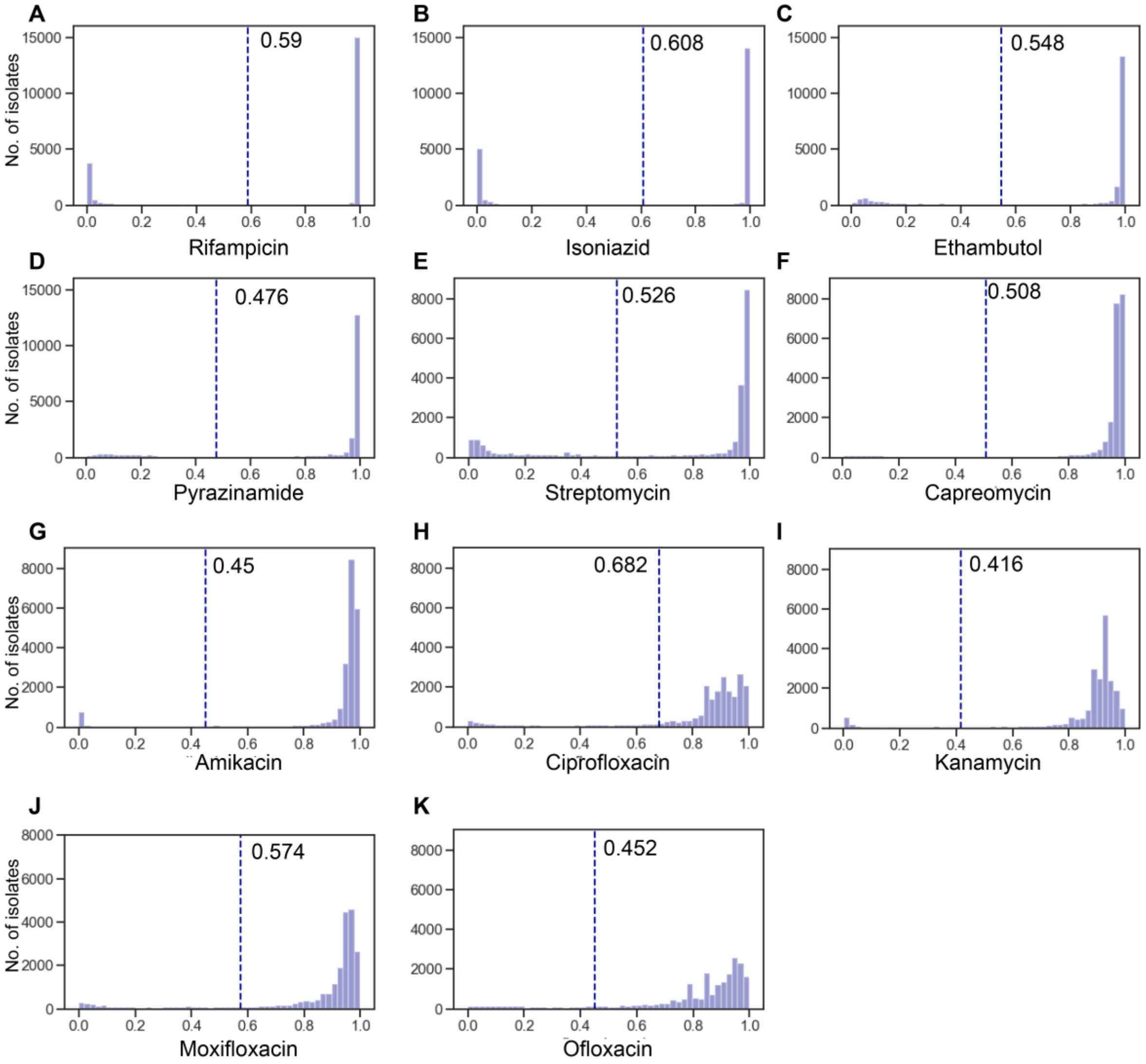


**Supplementary Figure S3: Probability distribution and thresholds according to** (Chen et al. 2019) **of Gentb-WDNN resistance predictions.** A-K. Probability of susceptibility (1 = drug susceptibility, 0 = drug resistance) for 11 drugs among the 20.379 isolates as predicted by Gentb-WDNN. Vertical lines depict the resistance threshold that yielded the highest predictive performance as measured by the sum of sensitivity and specificity. Drug-specific probability values are written in each subpanel.

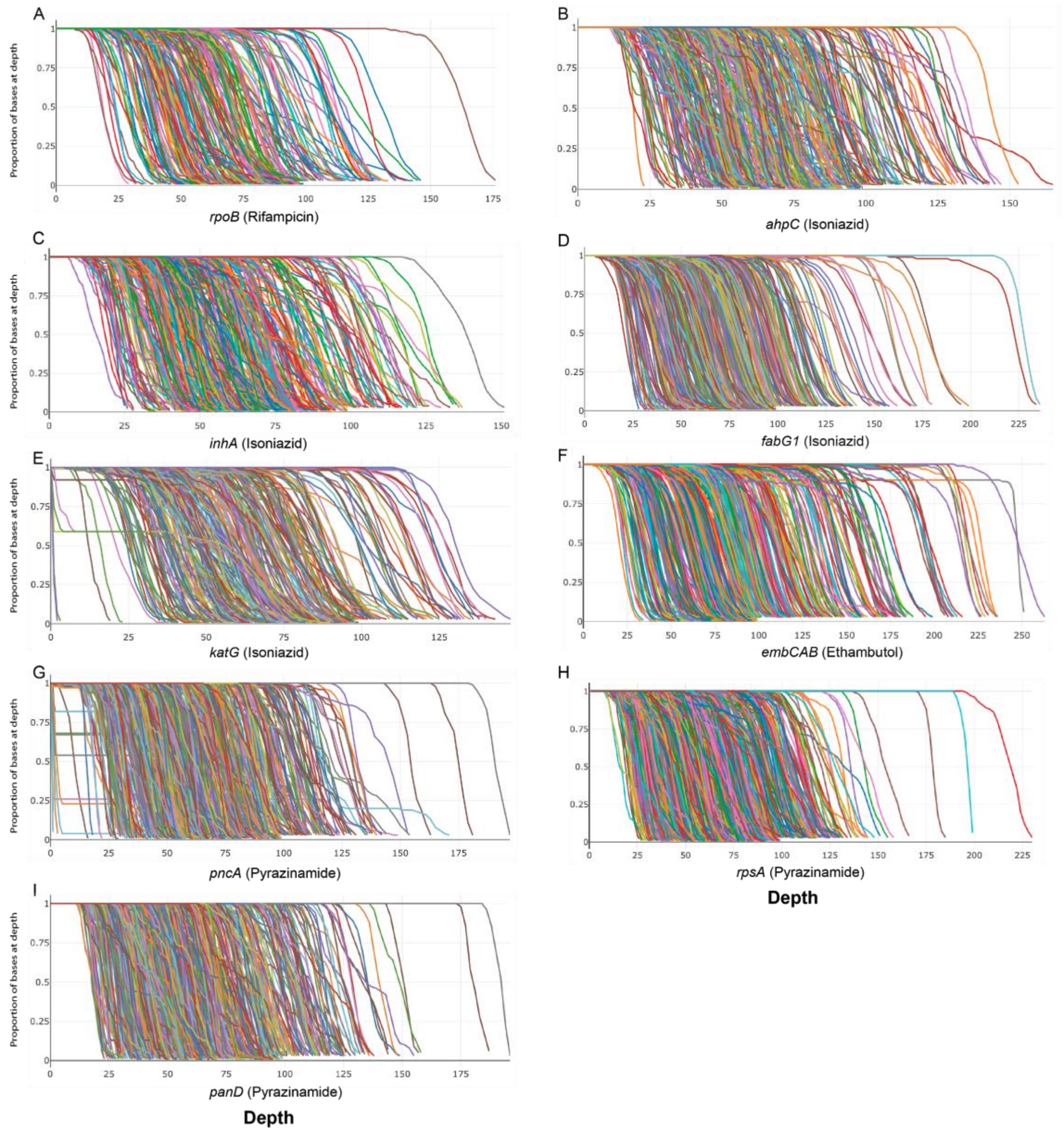


**Supplementary Figure S4: Sequencing depth of resistance conferring genes in isolates falsely predicted susceptible to first line agents.** The plots **A)** to **I)** display the sequencing depth against the proportions of bases covered at this depth for the length of the respective genes. Each line represents one isolate that was predicted susceptible by GenTB-RF while phenotypically resistant.

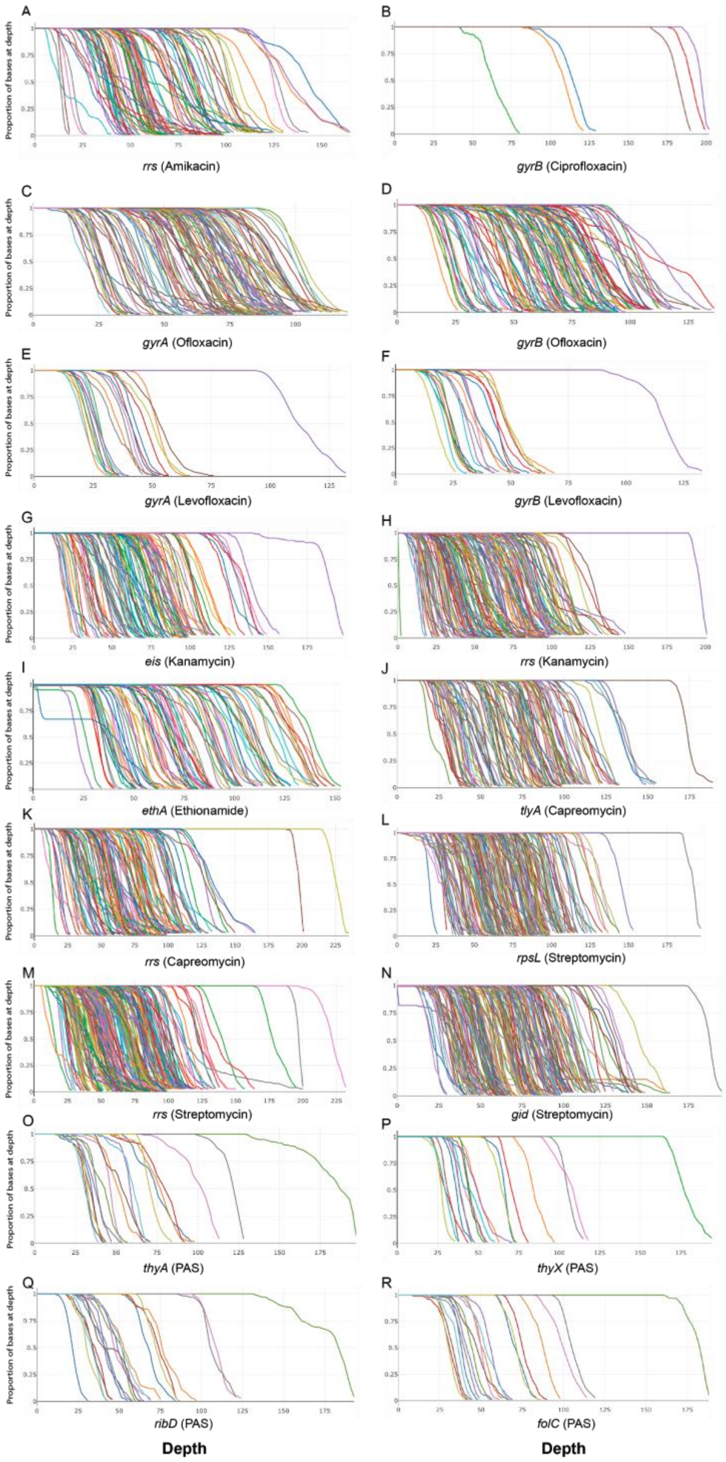


**Supplementary Figure S5: Sequencing depth of resistance conferring genes in isolates falsely predicted susceptible to second line agents.** The plots **A) to R)** display the sequencing depth against the proportions of bases covered at this depth for the length of the respective resistance genes noted below each plot. Each line represents one isolate that was predicted susceptible by GenTB-RF while phenotypically resistant.

**Supplementary Table S1:** Accessions and phenotype for sequence data used in this study
*[too large to put in here]*

**Supplemental Table S2:** Frequencies and percentages of available drug susceptibility data per drug

| Drug name | Resistant | | Susceptible | | Unknown | |
|---|---|---|---|---|---|---|
| | *n* | % | *n* | % | *n* | % |
| amikacin | 623 | 3.1 | 3,563 | 17.5 | 16,193 | 79.5 |
| capreomycin | 652 | 3.2 | 3,846 | 18.9 | 15,881 | 77.9 |
| ciprofloxacin | 63 | 0.3 | 331 | 1.6 | 19,985 | 98.1 |
| ethambutol | 3,001 | 14.7 | 12,788 | 62.8 | 4,590 | 22.5 |
| ethionamide | 502 | 2.5 | 1,095 | 5.4 | 18,782 | 92.2 |
| isoniazid | 6,141 | 30.1 | 13,509 | 66.3 | 729 | 3.6 |
| kanamycin | 583 | 2.9 | 3,878 | 19.0 | 15,918 | 78.1 |
| levofloxacin | 111 | 0.5 | 69 | 0.3 | 20,199 | 99.1 |
| moxifloxacin | 426 | 2.1 | 4,149 | 20.4 | 15,804 | 77.6 |
| ofloxacin | 762 | 3.7 | 4,313 | 21.2 | 15,304 | 75.1 |
| para.aminosalicylic_acid | 46 | 0.2 | 478 | 2.3 | 19,855 | 97.4 |
| pyrazinamide | 2,374 | 11.6 | 12,199 | 59.9 | 5,806 | 28.5 |
| rifampicin | 5,155 | 25.3 | 14,885 | 73.0 | 339 | 1.7 |
| streptomycin | 2,150 | 10.6 | 5,012 | 24.6 | 13,217 | 64.9 |
| **MDR & XDR** | | | | | | |
| MDR | 4,743 | 23.3 | - | - | - | - |
| XDR | 396 | 1.9 | - | - | - | - |

Note: MDR = Multi drug-resistant, XDR = Extensively drug-resistant.

**Supplementary Table S3:** Diagnostic accuracy comparison of tools for drugs with insufficient phenotype data and pyrazinamide performance on all isolates

| Drug | Phenotype | | GenTB - RF | | GenTB - WDNN | | Mykrobe | | TB-Profiler | |
|---|---|---|---|---|---|---|---|---|---|---|
| | | | Isolates sequenced with high depth (n = 19.880) | | | | | | | |
| | R (n) | S (n) | Sensitivity (95% CI) | Specificity (95% CI) | Sensitivity (95% CI) | Specificity (95% CI) | Sensitivity (95% CI) | Specificity (95% CI) | Sensitivity (95% CI) | Specificity (95% CI) |
| ciprofloxacin | 63 | 330 | 78% (66 to 88) | 98% (97 to 99) | 93% (85 to 100) | 97% (95 to 99) | 66% (53 to 78) | 98% (97 to 100) | 90% (83 to 97) | 98% (97 to 100) |
| levofloxacin | 65 | 104 | 81% (73 to 88) | 77% (66 - 87) | - | - | - | - | 74% (65 to 83) | 75% (64 to 86) |
| para-aminosalicylic_acid | 46 | 474 | 9% (2 to 18) | 100% (99 to 100) | - | - | - | - | 30% (17 to 44) | 98% (96 to 99) |
| pyrazinamide | 2,336 | 11,932 | 90% (88 to 91) | 88% (87 to 90) | 81% (79 to 82) | 95% (94 to 95) | 72% (71 to 74) | 98% (97 to 98) | 81% (80 to 83) | 96% (96 to 97) |

Note: Tool's performance on all isolates with available pyrazinamide phenotype shown, for performance on the hold-out validation dataset after Random Forest retraining please refer to Table 1.

**Supplementary Table S4** Area under the Receiver Operating Characteristic curve for GenTB-RF and GenTB-WDNN

| Drug | GenTB-RF | GenTB-WDNN |
|---|---|---|
| | Area under the ROC curve (95% CI) | |
| ciprofloxacin | 0.88 (0.82 to 0.93) | 0.95 (0.91 to 0.99) |
| levofloxacin | 0.79 (0.72 to 0.85) | - |
| para-aminosalicylic_acid | 0.54 (0.51 to 0.59) | - |

RF = Random Forest, WDNN = Wide and Deep Neural Network

**Supplementary Table S5:** Diagnostic accuracy to rifampicin and isoniazid across low-depth and passed-depth isolates.

| Tool | Drug | Low-depth isolates (n = 499) | | Passed-depth isolates (n = 19,880) | |
|---|---|---|---|---|---|
| | | Mean Sensitivity (SD) | Mean Specificity (SD) | Mean Sensitivity (SD) | Mean Specificity (SD) |
| GenTB-RF | isoniazid | 84.6 (3.64) | 98.2 (0.66) | 91 (0.36) | 97.6 (0.13) |
| GenTB-RF | rifampicin | 87.3 (3.64) | 98.5 (0.59) | 93.4 (0.37) | 98 (0.1) |
| GenTB-WDNN | isoniazid | 83.7 (3.77) | 99.4 (0.36) | 89.9 (0.4) | 98.9 (0.09) |
| GenTB-WDNN | rifampicin | 81.5 (4.19) | 99 (0.48) | 88.5 (0.45) | 98.9 (0.09) |
| TBProfiler | isoniazid | 75.6 (4.3) | 98.5 (0.61) | 91.1 (0.37) | 97.9 (0.13) |
| TBProfiler | rifampicin | 82.7 (4.05) | 98.8 (0.54) | 91.8 (0.40) | 98.3 (0.1) |
| Mykrobe | isoniazid | 70.4 (4.53) | 98.7 (0.56) | 86.7 (0.44) | 97.9 (0.13) |
| Mykrobe | rifampicin | 76.9 (4.49) | 98.7 (0.54) | 89.7 (0.44) | 98.5 (0.1) |

Note: GenTB-RF = GenTB Random Forest, GenTB-WDNN = GenTB - Wide and Deep Neural Network, SD = Standard Deviation

**Supplementary Table S6**: Non-silent variants in the gene *rpoB* among isolates with discordant phenotype and genotype predictions for the drug rifampicin

| False negative predictions by GenTB-RandomForest (*n* = 333 isolates) | | False positive predictions by GentTB-RandomForest (*n* = 254 isolates) | |
|---|---|---|---|
| Variant | count | variant | count |
| INS_CI_761103_i1296TTC_433F_rpoB[¶] | 14 | SNP_CN_761155_C1349T_S450L_rpoB[¶] | 49 |
| INS_CI_761135_i1328GAC_443L_rpoB[¶] | 9 | SNP_CN_761095_T1289C_L430P_rpoB[¶] | 33 |
| SNP_CN_761101_A1295T_Q432L_rpoB[¶] | 9 | SNP_CN_761139_C1333A_H445N_rpoB[¶] | 31 |
| DEL_CD_761101_d1294AATTCATGG_432_rpoB[¶] | 8 | SNP_CN_761889_G2083C_V695L_rpoB | 30 |
| DEL_CD_761115_d1308AAC_437_rpoB[¶] | 6 | SNP_CN_761109_G1303T_D435Y_rpoB[¶] | 30 |
| SNP_CN_760555_A749G_E250G_rpoB | 5 | SNP_CN_761277_A1471T_I491F_rpoB | 29 |
| DEL_CF_763258_d3451G_1151_rpoB | 4 | SNP_CN_761161_T1355C_L452P_rpoB[¶] | 26 |
| DEL_CD_761105_d1298CATGGA_433_rpoB[¶] | 3 | SNP_CN_761110_A1304T_D435V_rpoB[¶] | 9 |
| DEL_CD_761083_d1276GCACCA_426_rpoB[¶] | 2 | SNP_CN_761139_C1333T_H445Y_rpoB[¶] | 5 |
| DEL_CF_762516_d2709G_904_rpoB | 2 | SNP_CN_761155_C1349G_S450W_rpoB[¶] | 5 |
| INS_CI_761099_i1292CCA_431S_rpoB[¶] | 2 | SNP_CN_761139_C1333G_H445D_rpoB[¶] | 4 |
| SNP_CN_761141_C1335A_H445Q_rpoB[¶] | 2 | SNP_CN_761167_C1361T_P454L_rpoB | 3 |
| DEL_CD_761069_d1262CAAGGAGTTCTTCGGCAC_421_rpoB | 2 | SNP_CN_761110_A1304G_D435G_rpoB[¶] | 3 |
| DEL_CD_761100_d1293CAA_432_rpoB[¶] | 2 | SNP_CN_761140_A1334T_H445L_rpoB[¶] | 2 |
| DEL_CD_761088_d1281AGCCAGCTG_428_rpoB[¶] | 2 | SNP_CN_761880_G2074A_A692T_rpoB | 2 |

Note: the 15 most frequent variants or variant combinations are shown. Variants are denoted as follows: First type of variant, second if the variant leads to change in amino acid (AA), frameshift, or stop codon, third the genomic coordinate based on the reference strain H37RV (AL123456), fourth AA change, fifth codon change, last locus tag. [¶] Variant located in the rifampicin resistance determining region (RRDR)

**Supplementary Table S7**: Non-silent variants in the genes *inhA, katG, ahpC,* or *fabG1* among isolates with discordant phenotype and genotype predictions for the drug isoniazid

| False negative predictions by GenTB-RandomForest (*n* = 518 isolates) | | False positive predictions by GentTB-RandomForest (*n* = 315 isolates) | |
|---|---|---|---|
| Variant | count | Variant | count |
| SNP_CN_2155129_C983A_W328L_katG | 10 | SNP_CN_2155168_C944G_S315T_katG†¶ | 56 |
| SNP_CN_2154016_C2096T_G699E_katG | 6 | SNP_CN_1674481_T280G_S94A_inhA†¶ | 14 |
| SNP_CN_2155689_C423G_L141F_katG | 5 | SNP_CN_1674782_T581C_I194T_inhA† | 10 |
| SNP_CN_2155786_G326A_A109V_katG† | 5 | SNP_CN_2154695_C1417G_V473L_katG | 10 |
| SNP_CN_2154661_C1451T_R484H_katG | 4 | SNP_CN_2154075_C2037G_Q679H_katG | 2 |
| SNP_CN_2155665_C447G_W149C_katG | 3 | SNP_CZ_2154077_G2035A_Q679*_katG | 2 |
| SNP_CN_2155690_A422G_L141S_katG | 3 | SNP_CN_2726338_T146G_V49G_ahpC†¶ | 2 |
| SNP_CN_1674782_T581C_I194T_inhA† | 3 | SNP_CN_1674263_T62C_I21T_inhA† | 2 |
| SNP_CN_2155102_T1010C_Y337C_katG† | 3 | SNP_CN_2155168_C944T_S315N_katG†¶ | 2 |
| SNP_CN_1674262_A61G_I21V_inhA† | 3 | SNP_CN_2154676_G1436A_A479V_katG | 1 |
| SNP_CN_2154641_C1471T_G491S_katG | 3 | DEL_CF_2154510_d1602C_535_katG | 1 |
| SNP_CN_2154688_G1424A_T475I_katG | 3 | SNP_CN_2726323_C131G_P44R_ahpC† | 1 |
| SNP_CN_2155819_T293C_Y98C_katG | 3 | SNP_CN_2155258_C854G_G285A_katG | 1 |
| SNP_CN_2155222_C890A_G297V_katG | 3 | SNP_CN_2154760_C1352T_G451D_katG | 1 |
| SNP_CN_2154730_T1382G_Q461P_katG | 3 | SNP_CN_2155648_T464C_Y155C_katG† | 1 |

Note: The 15 most frequent variants are shown; Known lineage markers are excluded. † Variants that GenTB-RF has seen before. ¶ Variant considered important for isoniazid resistance by GenTB-Random Forest. We excluded variants in genes *kasA* and *embB* as their role in isoniazid resistance is questioned.
